# A Modularized Stretchable Microneedle Array for Minimally Invasive and Efficient Drug Delivery

**DOI:** 10.64898/2026.01.22.701215

**Authors:** Shuhuan Zhang, Yu Chang, Yuqing Fnu, Ke Du, Rui Liu

## Abstract

A novel Modularized Stretchable Microneedle Array (MSMA) is developed for minimally invasive and efficient drug delivery. By applying a modular design, the MSMA can be highly customized, offering versatility for different therapeutic applications. Its high stretchability allows for simultaneous stretching of both the microneedle and the skin, enhancing the penetration rate. Evaluations using tissue-mimicking materials and porcine skin demonstrated significant improvements in penetration efficiency and reduced residual strain. The MSMA achieved penetration rates of up to 89% on porcine skin under a minimum push-down distance of 2 mm. Moreover, the MSMA’s ability to stretch and release in synchronization with the skin significantly reduces residual strain. This innovative platform promises reduced pain and discomfort, representing a substantial advancement in transdermal drug delivery technology.

## Introduction

Microneedles, small-scale medical devices, are utilized for the administration of vaccines, drugs, and other therapeutic agents ^1^. In contrast to oral drug delivery, microneedles offer limited drug absorption or degradation by the gastrointestinal tract or liver ^2^. Unlike hypodermic needle delivery, microneedles avoid direct contact with nerve cells and vessels ^3^, thereby minimizing potential tissue damage compared to conventional hypodermic needles ^4,5^. Furthermore, microneedles require less training for application, making them more accessible for physicians ^6,7^. Additionally, compared to transdermal drug delivery, microneedles penetrate the stratum corneum, a barrier of dead corneocytes, leading to enhanced drug delivery efficiency ^8^.

While microneedles offer numerous benefits over conventional drug delivery methods, a significant hurdle remains in their efficiency of drug delivery ^9^. Penetrating the stratum corneum with a microneedle, due to its micro-scale size, presents considerable challenges. The underlying mechanism involves indenting the skin to create sufficient deformation, thereby facilitating skin penetration ^10^. Due to their diminutive size, microneedles often require considerable force to be effectively inserted into the skin to a depth that ensures an adequate penetration rate ^11^. Such a requirement in traditional microneedles poses several drawbacks, notably the increased risk of discomfort and pain for the patient due to the forceful application needed. Additionally, the need for substantial force to drive the microneedles into the skin may raise concerns regarding tissue damage, inflammation, and potential complications ^12^. Moreover, a substantial force causes the microneedle to bend, increasing the risk of fracturing ^13–15^.Achieving an optimal penetration rate is crucial for effective drug delivery, making the enhancement of this rate a focal point of ongoing research in the field of microneedle development.

One typical method for increasing penetration rates involves employing harder materials for microneedle fabrication and optimizing microneedle designs ^16^. Various materials, including polymers, silicon, and metals, have been explored for microneedle fabrication, yet their applicability is often limited by manufacturability constraints ^17^. Therefore, optimization of design parameters such as interspace between microneedles and individual microneedle geometry has been pursued to enhance penetration rates. Increasing the interspace between microneedles has been found to boost penetration rates ^18^, albeit at the cost of reduced microneedle density, which could limit drug delivery capacity. Consequently, considerable research has focused on optimizing the geometry of individual microneedles, with designs such as cones ^19^, arrows ^20^, bevel tips ^21^, and pyramids ^22^ being explored.

Khanna et al. ^10^ highlighted that the fundamental principle of microneedle penetration involves generating sufficient energy by applying pressure to the skin. When this energy surpasses the skin’s potential energy barrier, penetration occurs. Consequently, regardless of the needle’s geometry, the primary goal of optimization should be to enhance energy delivery to the skin. The most direct approach to achieving this goal is by sharpening the needle ^23,24^, as evidenced by numerous prior studies demonstrating the high penetration rates of sharper needles ^25,26^. However, sharper microneedles pose fabrication challenges and are prone to breakage ^18^, which can significantly reduce penetration rates if a microneedle fractures.

Another method employed to enhance penetration rates involves stretching the skin. In the studies conducted by Shrestha and Stoeber ^27^ and Verbaan et al. ^28^, the skin was stretched during the testing of microneedle performance to ensure stable and efficient penetration rates. Kim et al. ^29^ conducted a detailed investigation into how stretching the skin can improve the penetrability of a microneedle patch.

They examined the impact of skin stretching on the push-down distance required to break the skin surface. Their findings revealed that by uniaxially stretching the skin to an extension ratio of 1.2, the push-down distance could be decreased by 14.5%. Moreover, under equal biaxial tension conditions, stretching the skin in both directions to an extension ratio of 1.17 reduced the push-down distance by 53.3%. Additionally, stretching the skin increased the depth to which the microneedle could be inserted, fulfilling an important requirement for microneedle-mediated drug delivery.

The necessity for a microneedle to be applied to stretched skin directly correlates with the findings from prior research. Notably, the simultaneous stretching of both the needle and the skin is paramount for optimal penetration. Disregarding this crucial synchronization risks the microneedle being forcibly pushed out of the skin’s surface ^30^. Furthermore, the absence of concurrent stretching of the needle could disrupt the skin’s recovery process, resulting in residual strain that elevates the risk of damaging the skin tissue. This vital interplay between needle and skin stretching, as established in earlier study ^31^, underscores the significance of their simultaneous manipulation to achieve successful penetration without compromising the skin’s integrity. Consequently, this research emphasizes the need to develop a microneedle design capable of stretching in coordination with the skin. This microneedle’s design and application should guarantee that it can extend concurrently with the skin, promoting a synchronized stretching action that facilitates efficient and safe penetration through the skin.

To summarize, a stretchable microneedle enabled by high resolution 3D printing is developed in this study. While using this modularized stretchable microneedle array (MSMA), the skin and microneedle are stretched simultaneously, reaching a high penetration rate without being pushed down deeply by stretching the skin. In addition, since the microneedle and skin are stretched and released simultaneously, the residual strain on skin after releasing is decreased by over 90%. Our work introduces a novel microneedle platform characterized by minimal invasiveness and residual strain, which is crucial for reducing pain and distress in various biomedical applications.

### Design and fabrication of modularized stretchable microneedle array (MSMA)

To achieve the aforementioned features, the Modularized Stretchable Microneedle Array (MSMA) has been structured with three key components: silicone strips, handles, and microneedle columns (Figure 1.a), and the microneedle columns together with two handles are assembled and secured at both ends by silicone strips (Figure 1.b). This configuration enables the microneedle columns to be uniformly extended along with the silicone strips when the handles are pulled (Figure 1.c) and ensures that the expansion and contraction of the MSMA are synchronized with the patient’s skin stretching during the application of the microneedle(Figure 1.d).

**Figure 1.**
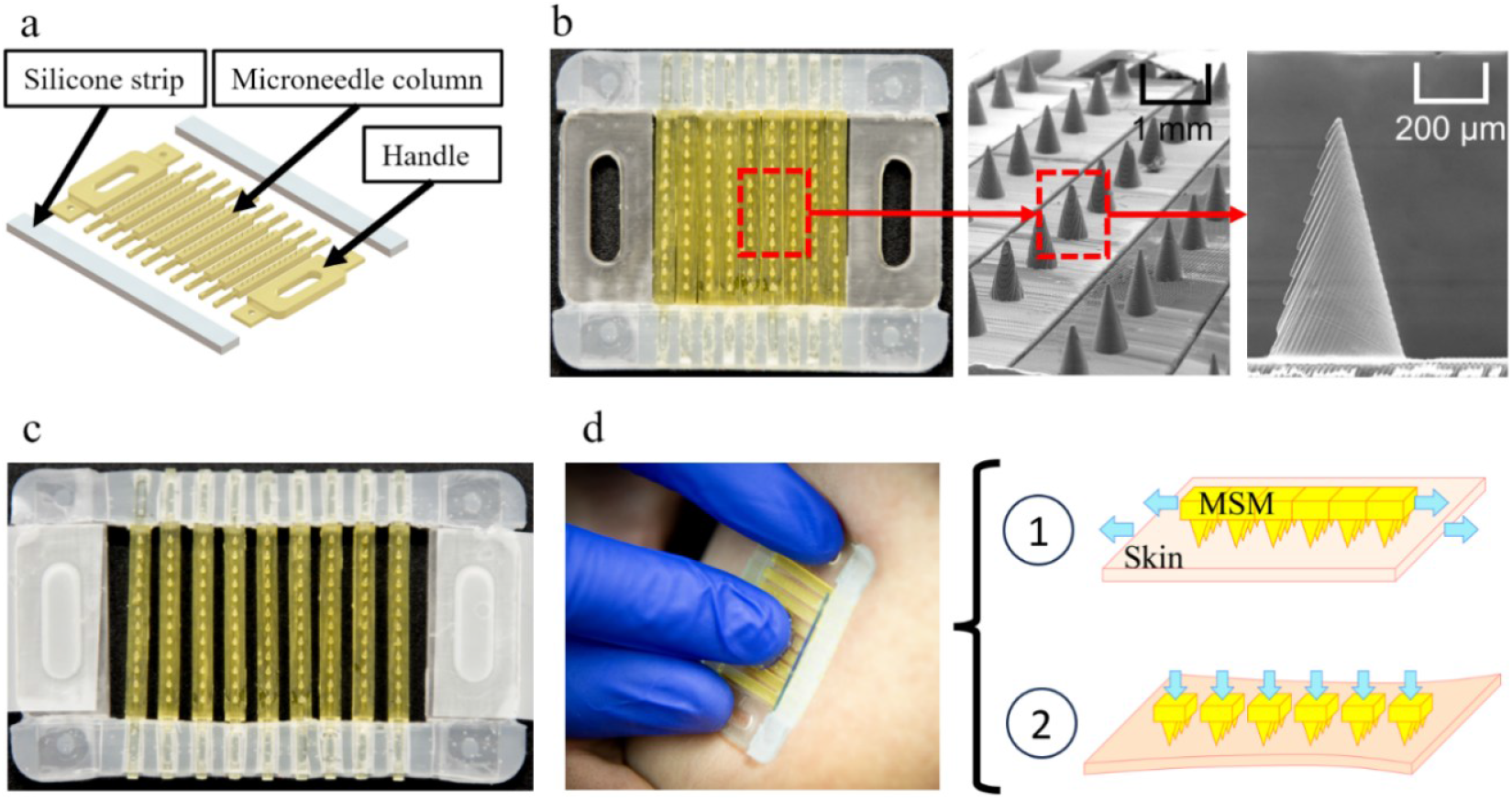
(a) Components of Modularized Stretchable Microneedle (MSM); (b) assembled MSM and the pictures of microneedles under TESCAN MIRA3 scanning electron microscopy (SEM); (c) stretched MSM; (d) two steps of operation process include simultaneously stretching the MSM and skin, then pushing downward the MSM to the skin.

The manufacturing procedure for MSMA is depicted in Figure 2. Initially, the microneedle columns and handles are created using HTL photosensitive resin with a microArch P140 3D printer, which ensures precise dimensions and robustness (Step 1) ^32,33^. In the next step, EcoFlex 00-50 silicone and a curing agent from Smooth-On are mixed in a 1:1 weight ratio and poured into the mold (Step 2). Nine microneedle columns, positioned between two handles, are then assembled and placed into the mold (Step 3). The EcoFlex 00-50 silicone is allowed to cure for three hours, after which the semi-finished microneedle structure is removed from the mold, with one end of the microneedle columns and handles secured in the silicone strip (Step 4). Following this, the same ratio of EcoFlex 00-50 silicone and curing agent is mixed and poured into the mold once more (Step 5). The semi-finished microneedle assembly is then flipped, and the other end of the microneedle columns and handles is carefully inserted into the mold (Steps 6 and 7). After another three-hour curing period and demolding, the MSMA is fully assembled and ready for use (Step 8).

**Figure 2.**
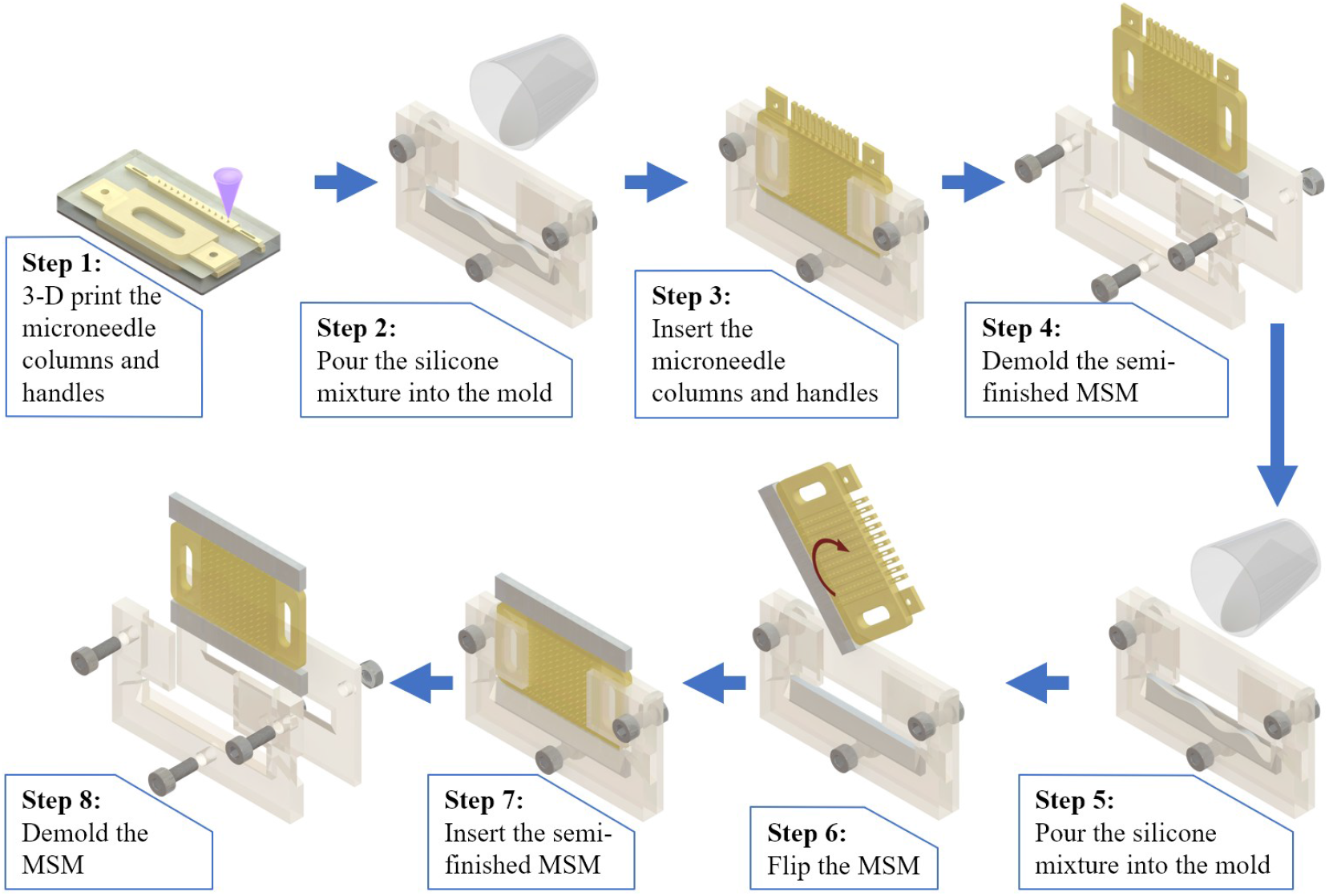
Initially, the microneedle columns and handles are created using HTL photosensitive resin with a microArch P140 3D printer, which ensures precise dimensions and robustness (Step 1). In the next step, EcoFlex 00-50 silicone and a curing agent from Smooth-On are mixed in a 1:1 weight ratio and poured into the mold (Step 2). Nine microneedle columns, positioned between two handles, are then assembled and placed into the mold (Step 3). The EcoFlex 00-50 silicone is allowed to cure for three hours, after which the semi-finished microneedle structure is removed from the mold, with one end of the microneedle columns and handles secured in the silicone strip (Step 4). Following this, the same ratio of EcoFlex 00-50 silicone and curing agent is mixed and poured into the mold once more (Step 5). The semi-finished microneedle assembly is then flipped, and the other end of the microneedle columns and handles is carefully inserted into the mold (Steps 6 and 7). After another three-hour curing period and demolding, the MSMA is fully assembled and ready for use (Step 8).

## Materials

In this study, the penetration performance of the developed MSMA has been systematically characterized using tissue-mimicking materials first and subsequently validated on biological tissues.

A tissue mimic membrane was utilized, which was fabricated using Polydimethylsiloxane (PDMS) from Dow Corning. PDMS has been previously employed in research to emulate human skin ^18,34^. Consequently, the PDMS film was employed as a skin mimic material in this study to demonstrate the proof-of-concept. The dimensions of the PDMS film were 70 mm × 50 mm × 1 mm (length × width × thickness).

The preparation of PDMS specimens involved using the Sylgard 184 Silicone Elastomer Kit, comprising a Silicone Elastomer Base and Curing Agent. The two components were mixed in a weight ratio of 20:1 and then poured into an aluminum mold. Subsequently, the mixture underwent a vacuum degassing process for 5 hours to eliminate any trapped air bubbles. The mixture was cured for 12 hours at 65°C. After the curing process, the solidified PDMS specimens were carefully removed from the mold and made ready for subsequent testing.

Given the similarities between human and porcine skin ^35,36^, a sample of porcine skin (Catalog number: NC1275387) was obtained from Fisher Scientific to evaluate the performance of microneedles. The non-sterile skin sample, which had been shaved and frozen, included both epidermal and dermal layers. The thickness of the porcine skin was 1.524 mm +/-0.254 mm.

### Evaluating the penetration rate on tissue mimic film and porcine skin

The experimental setup employed for this test is depicted in Figure 3.a. Central to this setup is a pair of grippers designed to firmly grasp both the tissue mimic membrane and the microneedle. One gripper, situated on the left side, remains stationary and is equipped with a hook mount. The opposite gripper on the right side is mobile, driven by a linear motor to move towards the right. Both grippers feature hooks for securely grasping and stretching the microneedle. This setup enables simultaneous stretching of both the tissue mimic film and the microneedle.

**Figure 3.**
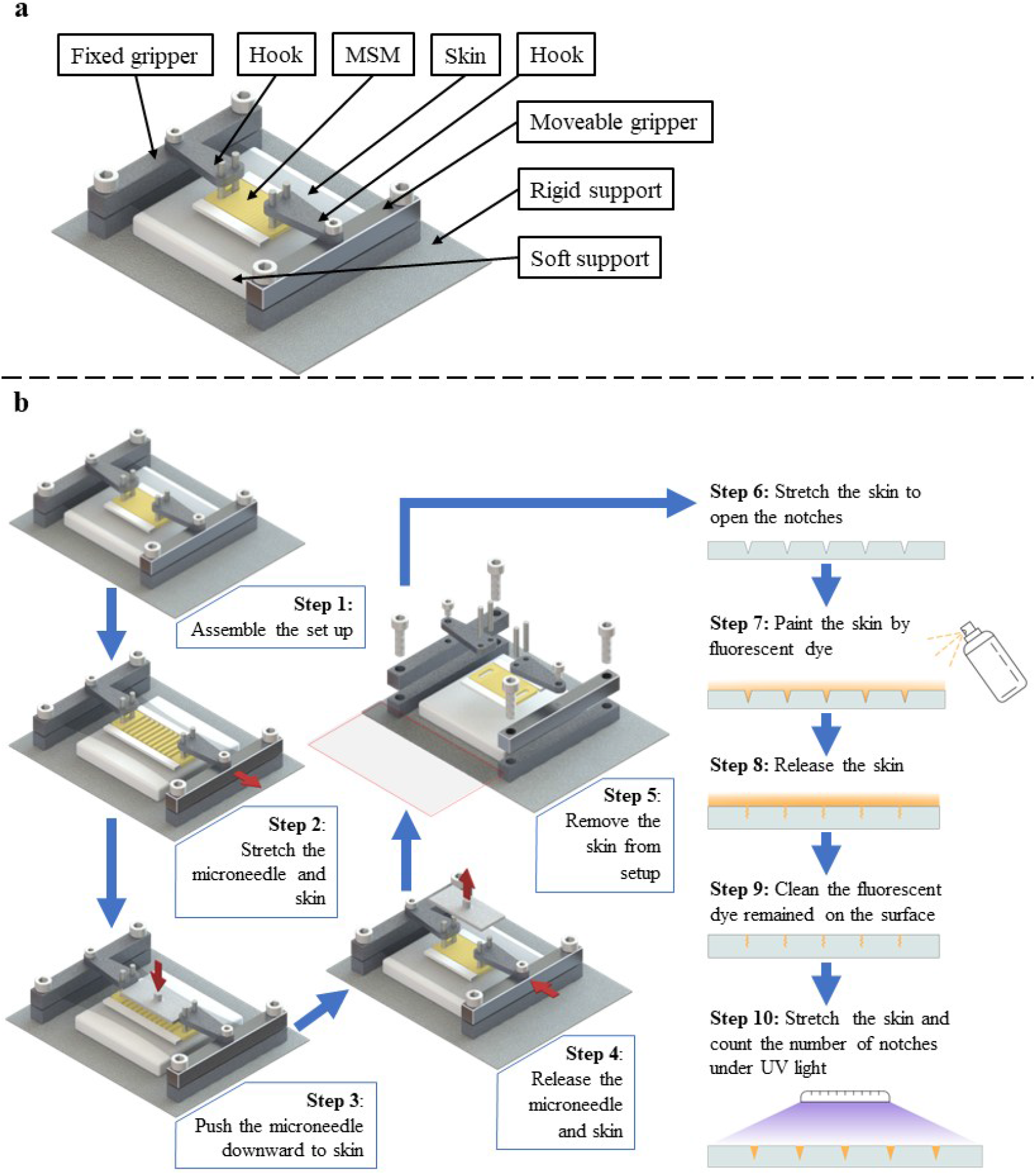
(a) Schematic drawing of the setup for testing the penetration rate of MSMA; (b) experimental process for testing the penetration rate of MSMA.

Additionally, the setup integrates a universal testing machine, which applies downward pressure onto the microneedle, facilitating its insertion into the tissue mimic film. To replicate the tissue and bone structures beneath the skin, a PDMS chunk is positioned underneath the tissue mimic membrane.

Firstly, the functionality of the developed MSMA was evaluated for its ability to penetrate the stretched sample, either tissue mimic materials or biological tissues, to a controlled depth using the experimental setup shown in Figure 3.b.

The procedure starts by securing a sample film between two grippers and attaching the microneedle handles (Step 1). This setup allows for the simultaneous stretching of the sample and the microneedle using a linear motor to achieve the predefined strain (Step 2). Following this, the microneedle is driven downward by a tensile tester to a predetermined distance and maintained in position for 4 seconds (Step 3). After this, the downward force is withdrawn, and both the sample and microneedle are released, with the microneedle being detached from the setup (Steps 4&5).

To accurately characterize the penetration rate from the microscale notches created by the microneedles, the sample undergoes an additional stretching (Step 6). Following this, a layer of fluorescent ink is applied to the sample’s surface and allowed to remain for 3 minutes, facilitating the infiltration of the ink into the notches (Step 7). The sample is then released (Step 8), and any superfluous fluorescent ink on the surface is removed (Step 9). Finally, the sample is stretched again and exposed to ultraviolet (UV) light, illuminating the fluorescent ink retained within the notches (Step 10). The penetration rate is calculated by dividing the number of illuminated notches by the number of microneedles.

The penetrating capability of the MSMA was initially evaluated on a tissue mimic film subjected to various tension strains and push-down distances. Tension strains of 0.05, 0.075, 0.1, 0.125, and 0.15 s were applied to the film. For each tension strain, the penetration rate was determined by depressing the microneedle to depths of 2, 3, and 4. To ensure accuracy and reliability, each combination of tension strain and push-down distance was replicated five times.

Subsequent evaluations of the MSMA’s penetration rate were conducted on porcine skin. Given the limited stretchability of porcine skin relative to living skin, the maximum tension strain was restricted to 0.1 to allow the skin to revert to its original state post-stretching. For comparative analysis, the penetration rate was also tested under a tension strain of 0.05, which corresponds to the minimum strain used on the tissue mimic film. In both scenarios, a push-down distance of 4 mm was applied to the porcine skin.

The penetration rates of the MSMA under various tension strains of the tissue mimic film and the corresponding images of the tissue mimic film under UV light are also presented in Figure 4. In Figure 4.a, it’s evident that, at a push-down distance of 2 mm, the penetration rate increases significantly from 31% to 87% with the rise in tension strain of the tissue mimic film from 0.05 to 0.15. When the push-down distances are increased to 3 mm and 4 mm, the effect of tension strain on penetration rate becomes less pronounced, which can be attributed to the high average penetration rates. Additionally, deeper push-down distances could cause skin discomfort. Therefore, compared to increasing the push-down distance, stretching the skin proves to be a more effective method for enhancing the penetration rate of microneedles. The details of the penetration rate under various tension strains and push-down distances are documented in Figure 4.b.

**Figure 4.**
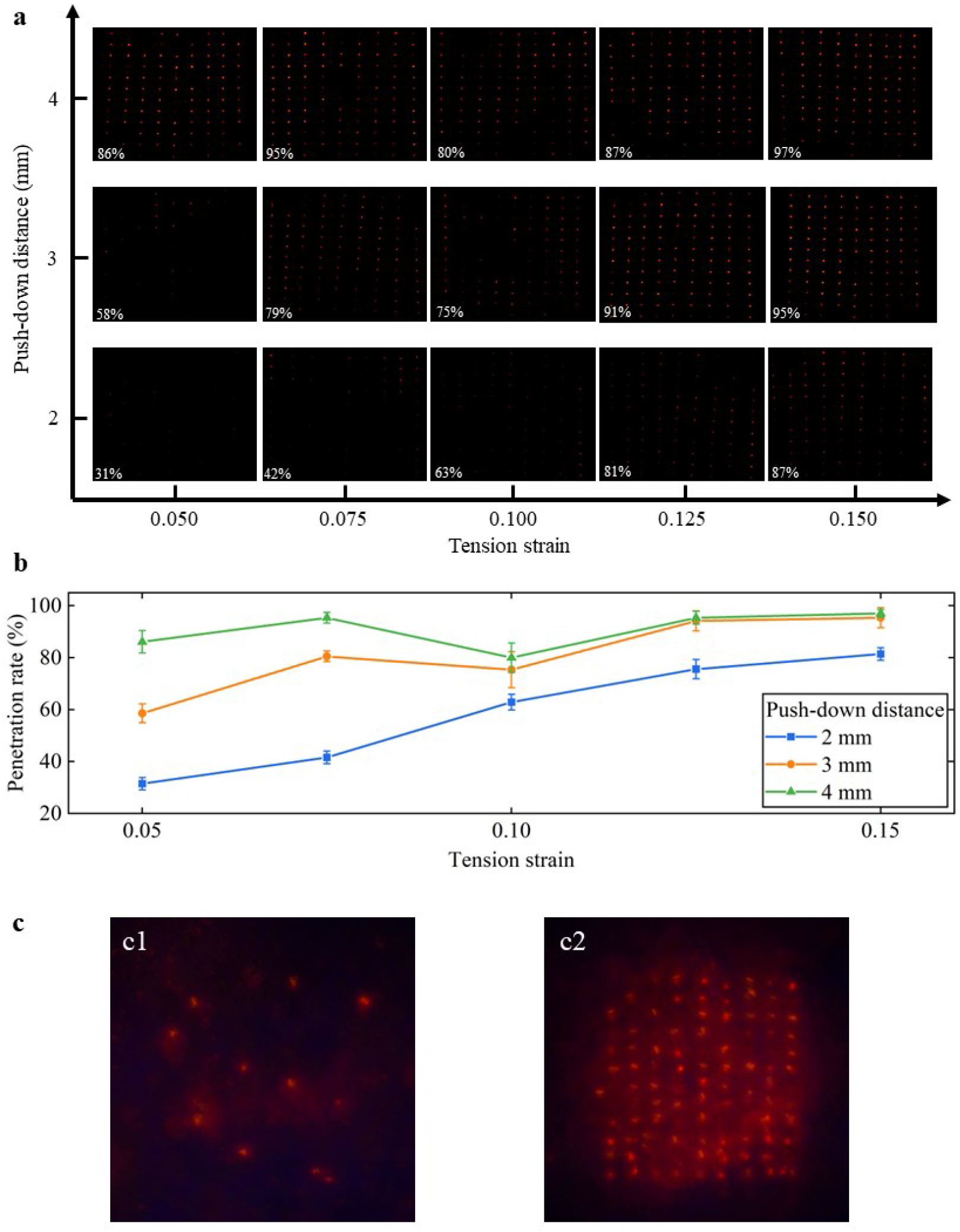
(a) Fluorescent marks under UV light demonstrate the penetration rate, displayed in the bottom left corner of each image, when the microneedle is pushed down for 2 mm, 3 mm, and 4 mm, under the tension strains of 0.05, 0.075, 0.1, 0.125, and 0.15. (b) The details of the penetration rate under various tension strains and push-down distances. (c, c1) The fluorescent marks on porcine skin under UV light demonstrate the penetration rate when the microneedle is pushed down for 4 mm and subjected to tension strains of 0.05. (c, c1) The fluorescent marks on porcine skin under UV light demonstrate the penetration rate when the microneedle is pushed down for 4 mm and subjected to tension strains of 0.1.

The penetration rate of the MSMA was further assessed on porcine skin. To ensure the porcine skin could return to its original state after stretching, a maximum tension strain of 0.1 was applied to the experimental group. For the control group, the penetration rate of the MSMA was tested under a tension strain of 0.05. In both groups, a push-down distance of 4 mm was applied to the porcine skin.

Under a tension strain of 0.05, only a few microneedles breached the surface of the porcine skin (Figure 4.c1), resulting in a penetration rate of 12%. However, by increasing the tension strain of the porcine skin to 0.1, a significantly higher number of microneedles penetrated the surface (Figure 4.c2), leading to a substantial increase in the penetration rate to 89%.

### Evaluating the residual strain on tissue mimic film and porcine skin

The setup utilized in this section is illustrated in Figure 5.a1. To offer a clearer depiction of the experimental components, a schematic drawing of the setup is provided in Figure 5.a2. The fundamental structure of this configuration closely resembles that of the setup used to evaluate the penetration rate. To quantify residual strain, a 3D Digital Image Correlation (DIC) system is employed. This DIC system consists of two cameras that track the movement of fluorescent markers on the sample. By analyzing the displacement of the fluorescent markers, the DIC system can compute the strain experienced by the sample.

**Figure 5.**
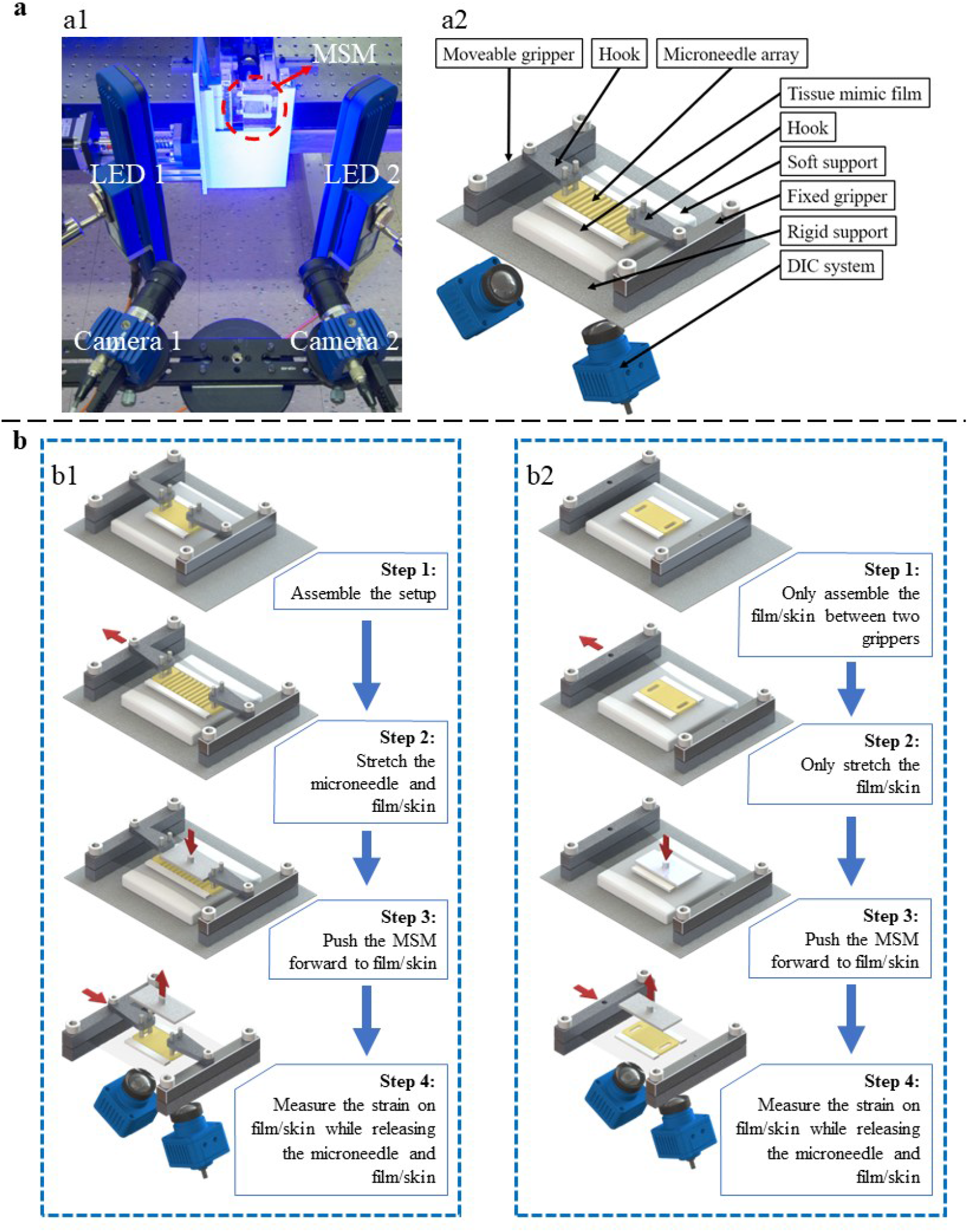
(a,a1) Experimental setup for investigating the residual strain. (a,a2) schematic drawing of the experimental setup for investigating the residual strain. (b,b1) experimental procedure of investigating the residual strain when stretching both the porcine skin and MSMA. (b,b2) experimental procedure of investigating the residual strain when only stretching the porcine skin.

To evaluate the hypothesis that significant residual strain persists on the sample when the sample and microneedle are not simultaneously stretched, two experimental scenarios were conducted to measure the residual strain on the tissue mimic film.

The influence of the proposed MSMA on sample retraction was investigated through the first scenario (illustrated in Figure 5.b1). The tissue mimic film was secured between two grippers, while the MSMA was held in place by two hooks. The sample film and MSMA were simultaneously stretched to a tension strain of 0.15, the maximum strain set in the previous experiment. Subsequently, the MSMA was depressed by 4 mm to achieve a high penetration rate, replicating conditions akin to actual medical applications. During the retraction process, the push-down force was released first, followed by the gradual release of the film and MSMA at the same speed. The residual strain stored in the tissue mimic film after complete retraction was measured using the DIC system.

In the comparative scenario (illustrated in Figure 5.b2) for the normal non-stretchable microneedle, only the tissue mimic film was clamped between the grippers, with the non-stretchable microneedle positioned atop the film. The film alone was stretched to a tension strain of 0.15. After stretching, the microneedle was depressed by 4 mm to penetrate the film. During the retraction process, the push-down force was released first, followed by the gradual release of the sample film. The residual strain resulting from the asynchronized retraction between the film and the non-stretchable microneedle was also measured.

The depiction of the residual strain on the tissue mimic film under two distinct testing conditions is displayed in Figure 6.a and Figure 6.b. Figure 6.a illustrates that the residual strain on the tissue mimic film is nearly imperceptible, signifying that most areas on the film can revert to their initial shape by undergoing simultaneous stretching and release with the MSMA. Conversely, when only the tissue mimic film is stretched without the simultaneous stretching of the MSMA (Figure 6.b), a conspicuous region (in red) with high strains (~0.15) appears in the middle of the film sample. This indicates that most of the applied strain during stretching is retained by the non-stretchable microneedle as residual strain, which can be attributed to the absence of simultaneous stretching and release.

**Figure 6.**
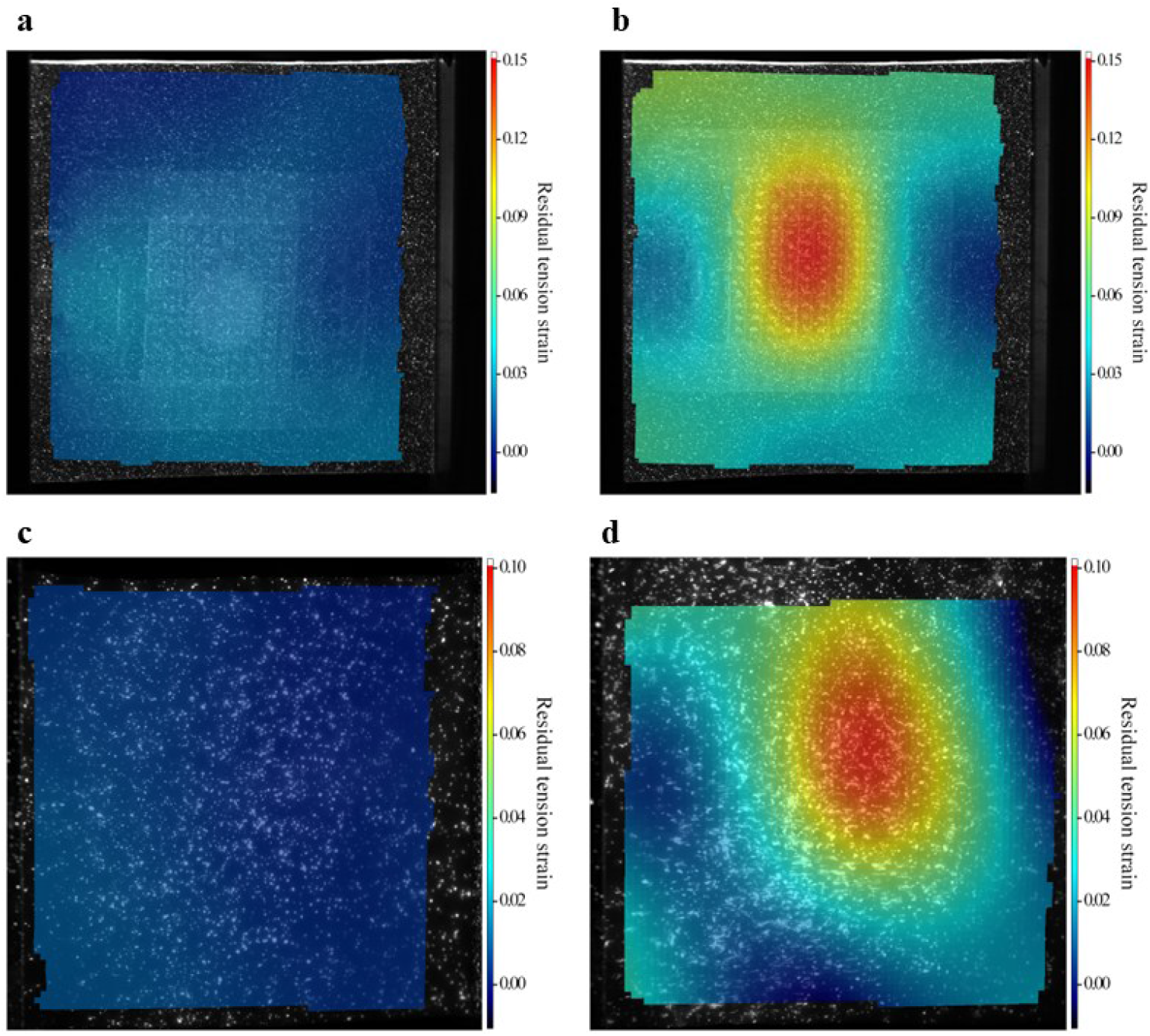
(a) Residual strain on the tissue mimic film after releasing the simultaneous stretch of both the tissue mimic film and MSMA; (b) residual strain on the tissue mimic film after releasing the stretch of tissue mimic film without MSMA; (c) Residual strain on the tissue mimic film after releasing the simultaneous stretch of both the porcine skin and MSMA; (d) residual strain on the tissue mimic film after releasing the stretch of porcine skin without MSMA.

The investigation also extended to evaluating the residual strain on porcine skin under two distinct operation processes of the MSN and the non-stretchable microneedle.

Given that the elasticity of the porcine skin sample may not be as robust as in vivo porcine skin, a tension strain of 0.1 is applied to the porcine skin in stretching to ensure that it can recover to its initial shape. All other testing steps and conditions remain consistent with those applied to the tissue mimic film.

When the MSMA and porcine skin were simultaneously stretched and released, the residual strain on the porcine skin was found to be uniform and close to zero (Figure 6.c). This suggests that the MSN has minimal impact on the recovery process of the porcine skin when stretched in conjunction with it. In contrast, when the stretched porcine skin was penetrated by the non-stretchable microneedle and then released, the microneedle impeded the skin’s recovery, resulting in significant residual strains (Figure 6.d). This indicates that the stretchability of the microneedle can significantly affect the skin’s ability to recover from stretching when not used in conjunction with it.

## Conclusions

The development of a modularized stretchable microneedle marks a promising advancement in transdermal drug delivery systems, addressing critical limitations of conventional microneedle designs ^37,38^. Its pioneering composition seamlessly integrates modularized microneedle columns, each individually fabricated, offering a unique opportunity to customize the size of the microneedle array. This pivotal feature allows tailoring the array to specific skin regions, ensuring precise and targeted drug delivery.

Moreover, the amalgamation of different microneedle column types—such as solid, hollow, and coated variants— within a single modularized microneedle imparts diverse functionalities. This adaptability empowers healthcare practitioners with unparalleled flexibility in selecting and administering drugs and therapies, enhancing treatment efficacy and personalization^39^.

Additionally, skin stretching during microneedle application plays a crucial role in the penetration rate. Intentional skin stretching facilitates microneedle penetration, significantly amplifying the penetration rate and thereby enhancing drug delivery efficiency. By leveraging skin mechanics alongside microneedle design, this technology overcomes traditional limitations related to push-down distance, ultimately improving drug delivery efficacy.

Another key advantage of the modularized stretchable microneedle system is its ability to synchronize the stretching of both the skin and the microneedle array. This synchronization supports controlled release mechanisms, allowing the microneedle array to retract seamlessly upon the release of the skin, without leaving any residual strain. This feature helps mitigate risks associated with residual strain, such as skin damage or microneedle dislodgment, thereby preserving both skin integrity and microneedle functionality. This minimally invasive procedure also provides faster healing, a lower risk of infection, and decreased inflammation, representing a major step forward in the advancement of microneedle technologies.

## ACKNOWLEDGEMENTS

Ke Du would like to thank for the support of NIH R35GM 142763

